# Olfactory sensory preconditioning in *Drosophila*: role of memory forgetting in gating S1/S2 associations

**DOI:** 10.1101/2021.11.29.470429

**Authors:** Anna Phan, Juan Martinez-Cervantes, Isaac Cervantes-Sandoval

**Author notes:** Correspondence: Isaac Cervantes-Sandoval, Department of Biology, Georgetown University, 37^th^ and O street, Washington, D.C.

## Abstract

Learning and memory storage is a complex process that has proven challenging to tackle. It is likely that, in real nature, the instructive value of reinforcing experiences is acquired rather than innate. The association between seemingly neutral stimuli increases the gamut of possibilities to create meaningful associations and increases the predictive power of moment-by-moment experiences. Here we report physiological and behavioral evidence of olfactory unimodal sensory preconditioning in fruit flies. We show that the presentation of a pair of odors (S1 and S2) before one of them (S1) is associated with electric shocks elicits a conditional response not only to the trained odor (S1) but to the odor previously paired with it (S2). This occurs even if the S2 odor was never presented in contiguity with the aversive stimulus. In addition, we show that inhibition of the small G protein and known forgetting regulator *Rac1* facilitates the association between S1/S2 odors. These results indicate that flies can infer value to non-paired odor based on the previous associative structure between odors, and inhibition of *Rac1* lengthens the time of olfactory “sensory buffer,” allowing linking of neutral odors presented in sequence.

## Introduction

Learning and memory are fundamental for animal survival in noisy and dynamic environments and for humans’ high cognitive abilities. A significant amount of what we know on the biological basis of learning and memory has come from studying pavlovian conditioning^1–7 8– 10 11–13^. In pavlovian conditioning, learning depends on the mere contiguity of conditioned and unconditioned stimuli. In *Drosophila*, pavlovian conditioning is mainly studied using classical olfactory conditioning. The mushroom bodies (MB) are the main brain structure involved in coding olfactory memories^11–13^. During olfactory memory acquisition, positive or negative values to initially neutral odors are assigned by reinforcement (electric foot shock or sugar reward). This reinforcement is achieved by the coincident activation of a sparse number of Kenyon cells (KC) by odorant and dopaminergic neurons (DAN) that innervate discrete zones, composed of 15 tile-like compartments of the MB lobes ^14–17^. Each of these tiles has a corresponding mushroom body output neuron (MBON), activation of which favors either approach or avoidance behavior ^18,19^. The molecular detection of the coincidence ^14,20,21^ is thought to change the output weight of KC synapses onto the corresponding MBON (KC>MBON), suggesting a model in which dopamine-induced plasticity tilts the overall MBON network to direct appropriate behavior ^19,22–28^. Studying classical conditioning has provided essential insights into the molecular, cellular, and circuit basis of how the brain transforms sensory information into memories and how it uses these memories to drive behavior. Nevertheless, in humans and other animals, the instructive value in real natural reinforcing experiences is acquired rather than innately instructive and does not dependent on mere contiguity. Learning theory has postulated the idea that learned behavioral control uses two types of information. The first, results from habits, policies, or cached values (*i*.*e*., pavlovian conditioning – model-free learning). This kind of information produces rapid, efficient behavior but does consider changes in the value of the expected outcome. The second type of information relies on the knowledge of the associative structure of the environment to infer value (model-based learning). In other words, acquiring new knowledge depends on creating an associative structure of external events to use it then to create additional new meaningful associations. Different types of high-order conditioning are examples of this type of information. In high-order conditioning procedures, neutral stimuli acquire the property to elicit conditional responses even though they are never in contiguity with a natural reinforcement. Second-order conditioning and sensory preconditioning are examples of high-order conditioning. In sensory preconditioning, two initially neutral stimuli (S1 and S2) are repeatedly presented in contiguity (preconditioning phase); later, one of the stimulus (S1) is paired with a reinforcer (conditioning phase). After this, S2 will elicit a conditioned response even when it was never paired with the reinforcer, indicating that preconditioning created an association between S1 and S2. The response to the preconditioned stimulus (S2) differs from the response to the reinforcer-paired S1 in that it is not based on a cached value; instead, it must reflect the subject’s ability to infer value by virtue of knowledge of the associative structure of the task ^29^.

A wealth of growing data has demonstrated that in mammals, the retrosplenial cortex (RSC) makes important contributions to forming these S1-S2 associations ^30–34^. This is consistent with the circuit architecture of the RSC. The RSC contains strong reciprocal connections to subcortical thalamic nuclei ^35–42^. It also contains reciprocal connections with multiple sensory cortical regions and structures that compose the hippocampal and parahippocampal memory systems ^35,43–47^. Nevertheless, we still have a long way to go to better understand the neurobiological basis of sensory preconditioning. Thus, studies in anatomically simplified models, like *Drosophila*, –where behavioral effects can often be attributed to the actions of individual neurons ^18,48–51^– serve as high-resolution systems that can provide experimental flexibility and conceptual clarity. We suggest that understanding how a simple brain achieves sensory preconditioning with limited neurons and synapses will provide meaningful insights into how a more complex mammalian brain achieves the same task.

High-order conditioning has been reported in honey bees and *Drosophila*^52,53^. Using a fly simulator arena ^53^, reported the presence of both sensory preconditioning and second-order conditioning. Here we report evidence of olfactory unimodal sensory preconditioning in fruit flies. We show that the presentation of a pair of odors (S1 and S2) before one of them (S1) is associated with electric shocks elicits a conditional response not only to the trained odor (S1) but to the odor previously paired with it (S2). This occurs even if the S2 odor was never presented in contiguity with the negative reinforcer. These data provide evidence for unimodal sensory preconditioning in *Drosophila*. These findings open the door to better understand how a simple brain can achieve these forms of associations and will provide significant insights into how more complex brains solve similar problems.

## Results and Discussion

Using electrophysiology and vivo calcium imaging, it has been shown that pairing and odor with electric shocks induced a decreased response to the CS+ of MBON-γ1pedc>α/β ^26,54,55^. We recently reported a non-specific depression to non-associated odor observed when flies expressed the dominant-negative *Rac*^*N17*^ (*RacDN*) in KC compared to control animals ^55^. We recorded olfactory responses in MBON-γ1pedc>α/β of control animals exposed to single 5 s pulses of MCH (S1) followed by OCT (S2) with thirty seconds inter-stimulus interval (ISI). After, animals were trained by pavlovian conditioning by presenting a 20 s MCH pulse along with four 90 V, 1.25 s electric shocks. Post-training responses to MCH and OCT were recorded five minutes after conditioning. We observe a significant depression to OCT (S2) compared to the pre-response (Figure 1B). Flies kept at 18 C to keep target system at non-permissive temperature showed normal learning-induced odor depression to trained odor and no change to S2 ^55^. These results suggested that the pre-presentation was causing increased habituation and depression of the non-associated odor when *Rac1* is inhibited. To test this, we recorded olfactory responses in flies exposed to the same odor protocol except that the unconditioned stimulus was excluded. Results showed no depression to either odor presented (data not shown). Going even further, we confirm this observation by recording and evaluating changes in calcium responses after ten repeated presentations of both OCT and MCH. Data were analyzed similarly to previously describe^51^. The ratio between the mean of three initial responses and mean of three last responses in MBON-γ1pedc>α/β was not significantly different between control and experimental genotypes (data not shown).

**Figure 1.**
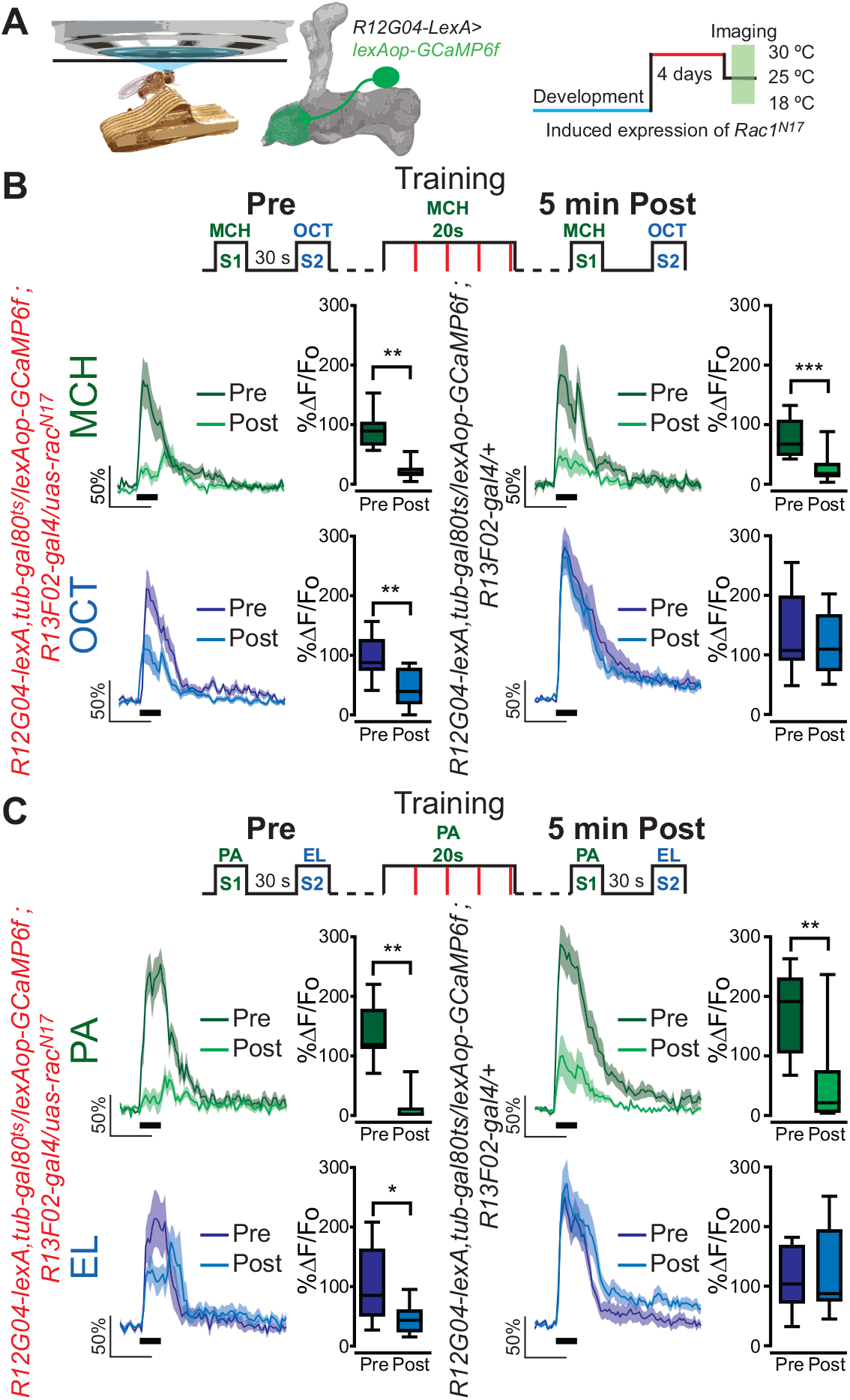
Rac1 inhibition induces depression to no trained pre-paired odor S2 in MBON-γ1pedc>α/β. (A) Diagram of in vivo under the microscope training, imaging setup, MBON-γ1pedc>α/β diagram, and *Rac1*^*N17*^ expression induction. (B) Upper part, diagram of experimental setup (odor schedule): preconditioning responses were obtained for MCH and OCT (S1 and S2); later flies were aversively trained to MCH (S1) and 5 min later post-conditioning responses were recorded. S1 (MCH) responses were completely depressed 5 min after training in both control and flies expressing *Rac1*^*N17*^. Non-parametric Wilcoxon-paired test *p≤0*.*0003*; *n=10-11*. No significant changes were detected to S2 (OCT) for control animals Non-parametric Wilcoxon-paired test *p=0*.*2783*; *n=11*. Nevertheless, a significant inhibition of post-trained response to S2 (OCT) was observed in flies expressing *Rac1*^*N17*^. Non-parametric Wilcoxon-paired test *p≤0*.*0098*; *n=10*. (C) Similar results were obtained for PA and EL. Upper part, diagram of experimental setup (odor schedule): preconditioning responses were obtained for PA and EL (S1 and S2); later flies were aversively trained to PA (S1) and 5 min later post-conditioning responses were recorded. S1 (PA) responses were completely depressed 5 min after training in both control and flies expressing *Rac1*^*N17*^. Non-parametric Wilcoxon-paired test *p≤0*.*0078*; *n=8-11*. No significant changes were detected to S2 (EL) for control animals Non-parametric Wilcoxon-paired test *p=0*.*5771*; *n=11*. Nevertheless, a significant inhibition of post-trained response to S2 (EL) was observed in flies expressing *Rac1*^*N17*^. Non-parametric Wilcoxon-paired test *p=0*.*0391*; *n=8*. Boxplots represent distribution of %ΔF/Fo responses across the 5 s of odor presentation. The thick black bar below each trace represents the time of odor presentation.

*Rac1* has been implicated in memory forgetting, and it has been theorized that forgetting is fundamental for memory generalization. In addition, Hige et al. ^54^ showed that this non-specific depression could be explained by partial overlapping in the representation of odors like OCT and MCH. This suggested that the expression of *Rac*^*N17*^ in KC was somehow increasing that generalization of MCH and OCT. To test this, we trained animals using more orthogonal odors, pentyl acetate (PA) and ethyl lactate (EL). Surprisingly, similar results to OCT/MCH were obtained; we observed a non-specific depression to the non-paired odor (EL) when flies expressed *Rac*^*N17*^ in KC (Figure 1C). We hypothesized that this non-specific depression could arise after aversive olfactory conditioning induces a memory generalization to any odor when *Rac*^*N17*^ is expressed. To test the first, we pre-presented OCT and MCH as before, trained the flies with MCH (S1), and then recorded four post-training odor responses (MCH, OCT, PA, and EL). This allowed us the possibility to evaluate changes to trained odor (MCH) (S1), a non-shocked paired previously experienced odor (OCT) (S2) and new never experienced odors (NO) (PA and EL). We compared these odor responses to responses of animals trained with backward conditioning. Surprisingly, expressing *Rac*^*N17*^ in KC induced a significant depression to OCT (S2) but left intact responses to novel odors, PA and EL (data not shown). To test this even further, we trained animals with forward or backward conditioning but excluding any pre-training odor presentation. Results showed significant depression to the odor associated with the shock but not any additional odors (data not shown). These results showed that the non-specific depression was not due to general odor habituation or generalization of learned aversion. Instead, depression of the non-paired odor is dependent on learning and the pre-presentation.

It has been reported that animals can link neutral sensory cues presented in succession or simultaneously; if then one of these cues is associated with reinforcement, animals can display memory to the non-associated pre-linked cue; this is called sensory preconditioning^29,56^. This kind of behavior is categorized as model-based learning where a value is inferred from knowledge of the associative structure of the environment^29^. Sensory preconditioning has been reported in honeybees^57^. In addition, it has been reported that expressing *Rac1*^*N17*^ in KC significantly enhances trace conditioning, in which an odor is associated with an electric shock presented many seconds after odor offset–suggesting that inhibition of *Rac1* lengthens an olfactory “sensory buffer” that later converges with the punishment signal ^58^. Therefore, we reasoned that sensory preconditioning and Rac1 inhibition could explain our results. To test this, we recorded olfactory responses in MBON-γ1pedc>α/β of wildtype flies and flies expressing *Rac1*^*N17*^ exposed to single pulses of MCH (S1) followed by OCT (S2) with a one-second inter-stimulus interval (ISI). Post-training responses to MCH, OCT, PA, and EL were recorded five minutes after conditioning. Using this reduced ISI, we observed a complete depression to the MCH (S1) and a significant reduction to OCT (S2) in wildtype animals when compared to animals trained with backward conditioning (data not shown). Responses to novel odors, PA and EL, showed no change. These results suggested that an association between neutral odors OCT/MCH (S1/S2) occurs during the pre-presentation and *Rac1* inhibition in KC extends the period for this association to occur. If this is true, we predicted that extending the ISI should eliminate sensory preconditioning in *Rac1DN* flies. As expected, no sensory preconditioning was observed in flies expressing *Rac1*^*N17*^ in KC when ISI was increased to 5 min (Figure 2B). These results indicate that flies show, at least at the physiological level, sensory preconditioning.

**Figure 2.**
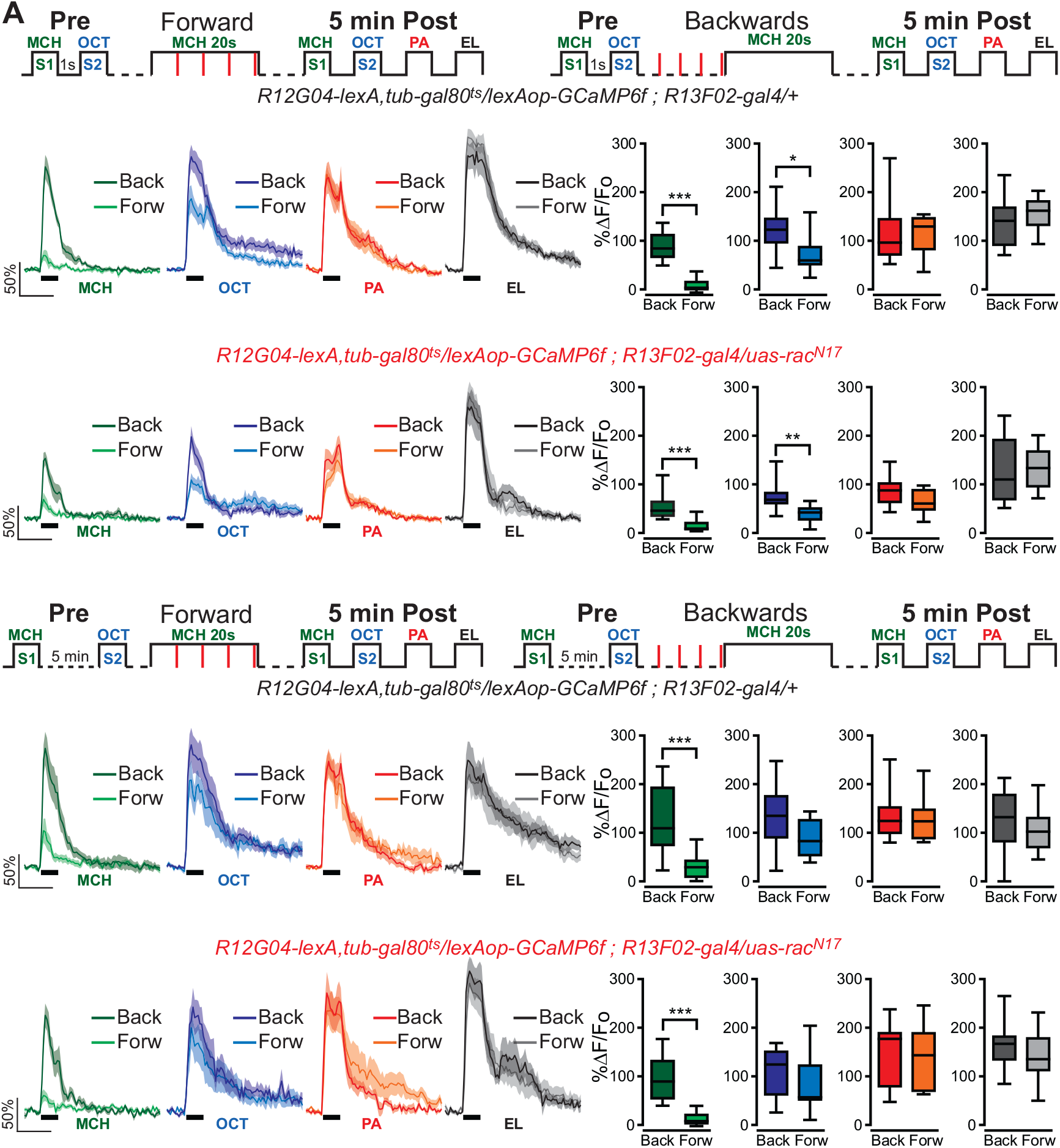
One second ISI during preconditioning induces depression to no trained pre-paired odor S2 in wildtype animals. (A) Upper part, diagram of experimental setup (odor schedule): preconditioning was induced by presentation of MCH and OCT (S1 and S2) with one second ISI; later flies were aversively trained to MCH (S1) and 5 min later post-conditioning responses were recorded. Responses were compared to flies trained using backwards conditioning. S1 (MCH) responses were completely depressed 5 min after training in both control and flies expressing *Rac1*^*N17*^. Non-parametric Mann-Whitney test *p≤0*.*0009*; *n=8-10*. A significant inhibition of post-trained response to S2 (OCT) was also present in control flies and flies expressing *Rac1*^*N17*^. Non-parametric Mann-Whitney test *p=0*.*035 and p=0*.*0031* respectively; *n=8-10*. Neither PA nor EL responses showed any significant reduction for both control and experimental flies. Non-parametric Mann-Whitney test *p≥0*.*0726, n=8-10*. (B) Upper part, diagram of experimental setup (odor schedule): preconditioning was induced by presentation MCH and OCT (S1 and S2) with 5 min ISI; later flies were aversively trained to MCH (S1) and 5 min later post-conditioning responses were recorded. Responses were compared to flies trained using backwards conditioning. S1 (MCH) responses were completely depressed 5 min after training in both control and flies expressing *Rac1*^*N17*^. Non-parametric Mann-Whitney test *p≤0*.*0044*; *n=8-10*. Neither OCT, PA, nor EL responses showed any significant reduction for both control and experimental flies. Non-parametric Mann-Whitney test *p≥0*.*1220, n=8-10*. Boxplots represent distribution of %ΔF/Fo responses across the 5 s of odor presentation. The thick black bar below each trace represents the time of odor presentation.

We then tested if sensory preconditioning was expressed at a behavioral level. For this, we trained the flies using an Arduino microcontroller for a precise odor delivery by controlling solenoids automatically. Using this Arduino system, we trained animals as follows. Wildtype flies were presented to a single pairing of the odors (5s pulse each) with a 1s ISI. Flies were then conditioned by the presentation of 1 min S1 along 12, 90 V shocks. Memory was tested right after training on a T-maze using either the S1 (shock-paired odor) and a novel never experience odor (NO) or S2 (non-shocked pre-paired odor) and a NO. As in canonical classical conditioning experiments, each behavioral experiment was conducted using the reciprocal odor as S1. The final performance index (PI) was calculated by averaging the PI for each odor used as S1. Results were compared to flies trained with backwards conditioning. Despite observing evidence of sensory preconditioning by functional imaging using a similar protocol, we could not observe any behavioral memory to the non-shocked pre-paired odor (S2) (Figure 3B). At a behavioral level, typically, sensory preconditioning is induced in animals by a repeated presentation of a pair of sensory cues. Preconditioning flies with ten repeated presentations of odor pairings (S1/S2) before training resulted in significant aversive memory expression to S2 (Figure 3B). These results suggest that a single odors pairing is sufficient to induce sensory preconditioned-related plasticity in MBON-γ1pedc>α/β compartment but is not enough to drive the learned behavior by S2 odor. We suggest that the presentation of additional odor pairings recruits and induces sensory preconditioning related plasticity in additional MB compartments and is the additive effect of multiple compartments that can then drive the behavior. Similar phenomena have been observed before in for classical conditioning where a short pairing of odor and shock – one second odor paired with 4 DAN photostimulation pulses – is enough to induce plasticity in MBON-γ1pedc>α/β compartment but in MBON-α2sc. Additional training – one minute odor presentation paired with 120 DAN photostimulation pulses – induces plasticity in MBON-α2sc ^54^. Preconditioning flies with MCH/OCT pairs repeated ten times before training resulted in a more robust depression in MBON-γ1pedc>α/β to the non-shocked pre-paired odor (CS2) in control flies (Figure 3B). These results were similar when PA/EL were used as the preconditioned odors (Figure 3C). Finally, we tested whether the expression of *Rac1*^*N17*^ in KC would increase the ISI at the behavioral level. Thirty seconds ISI resulted in no aversive memory to S2 in control flies but resulted in significant aversive memory in flies expressing *Rac1*^*N17*^ (Figure 4B). Flies kept at 18 C to keep target system at non-permissive temperature did not show evidence of sensory preconditioning (data not shown). Finally, as expected, increasing the ISI to 5 min restricted sensory preconditioning in flies expressing *Rac1*^*N17*^ (Figure 4B).

**Figure 3.**
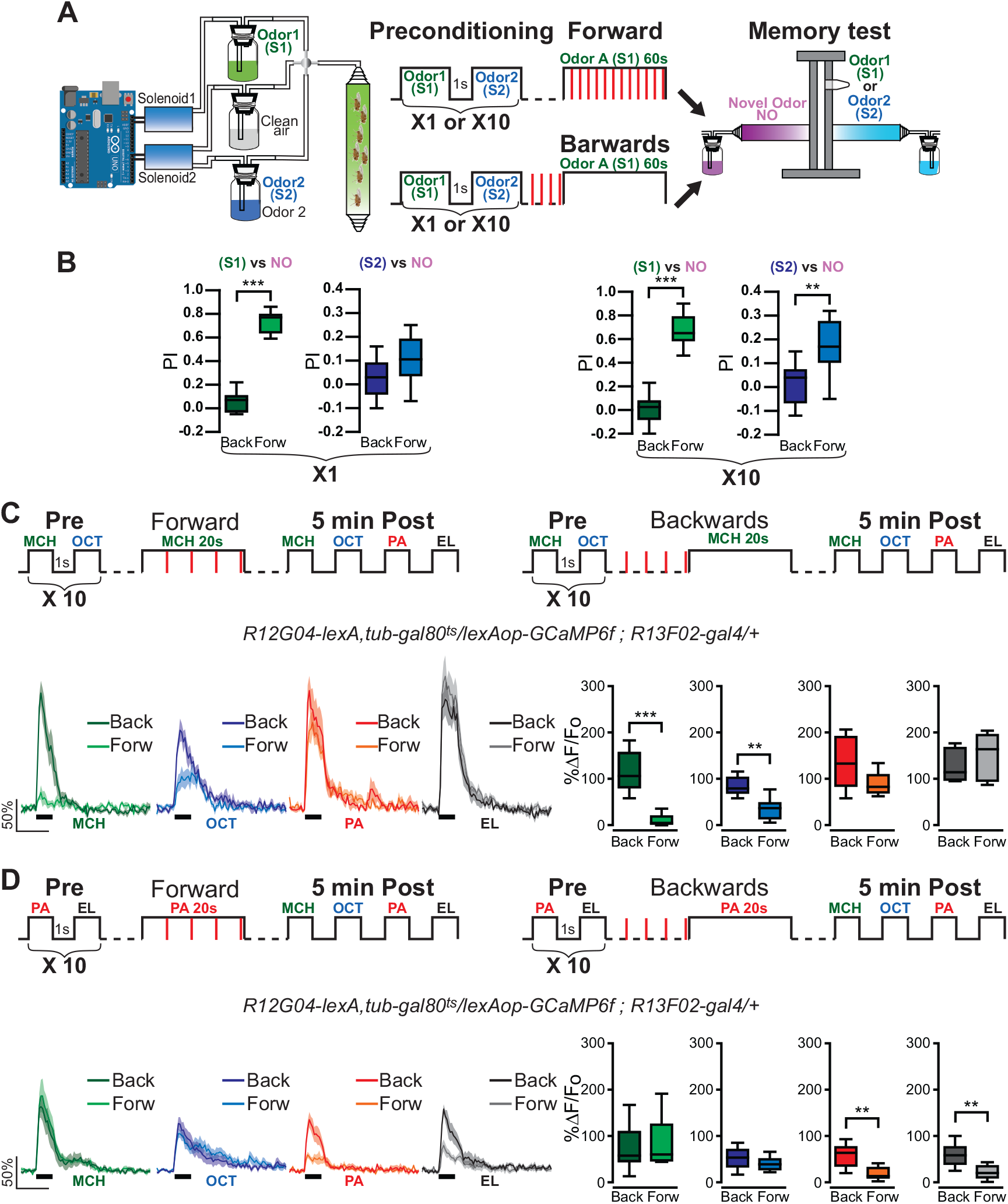
Repeated presentations of paired odors (S1/S2) induces behavioral sensory preconditioning. (A) Left, wildtype flies were trained using an Arduino microcontroller for precise odor delivery. Right, diagram of experimental setup (odor schedule): Flies were preconditioned by a single or ten repeated presentations of S1/S2 odor pairs with one second ISI. Later flies were aversively trained to MCH (S1) by pairing one minute odor presentation along 12 90V, 1.25 s shocks. Right after training memory was tested in a T-maze by presenting either S1 vs a novel odor (NO) or S2 vs a NO. Performance index (PI) were compared to flies trained using backwards conditioning. B) Memory to S1 was significantly different from flies trained with backwards conditioning, with both 1X or 10X preconditioning presentations. Non-parametric Mann-Whitney test *p<0*.*0001, n=9-12*. Memory to S2 was not significantly different from flies trained with backwards conditioning=, with 1X preconditioning presentations. Non-parametric Mann-Whitney test *p=0*.*0923, n=10*. In contrast, Memory to S2 was significantly different from flies trained with backwards conditioning, with 10X preconditioning presentations. Non-parametric Mann-Whitney test *p=0*.*0019, n=12*. C) Upper part, diagram of experimental setup (odor schedule): preconditioning was induced by ten repeated presentations of MCH and OCT (S1 and S2) with one second ISI; later flies were aversively trained to MCH (S1) and 5 min later post-conditioning responses were recorded for MCH, OCT, PA and EL. Responses were compared to flies trained using backwards conditioning. S1 (MCH) responses were completely depressed 5 min after training. Non-parametric Mann-Whitney test *p≤0*.*0043*; *n=6*. A significant inhibition of post-trained response to S2 (OCT) was also observed. Non-parametric Mann-Whitney test *p=0*.*0087*; *n=6*. Neither PA nor EL responses showed any significant reduction. Non-parametric Mann-Whitney test *p≥0*.*2468, n=6*. D) Upper part, diagram of experimental setup (odor schedule): preconditioning was induced by ten repeated presentations of PA and EL (S1 and S2) with one second ISI; later flies were aversively trained to PA (S1) and 5 min later post-conditioning responses were recorded for MCH, OCT, PA and EL. Responses were compared to flies trained using backwards conditioning. S1 (PA) responses were completely depressed 5 min after training. Non-parametric Mann-Whitney test *p≤0*.*0077*; *n=8*. A significant inhibition of post-trained response to S2 (EL) was also observed. Non-parametric Mann-Whitney test *p=0*.*0019*; *n=8*. Neither MCH nor OCT responses showed any significant reduction. Non-parametric Mann-Whitney test *p≥0*.*5054, n=8*. Boxplots represent distribution of %ΔF/Fo across the 5 s of odor presentation. The thick black bar below each trace represents the time of odor presentation.

**Figure 4.**
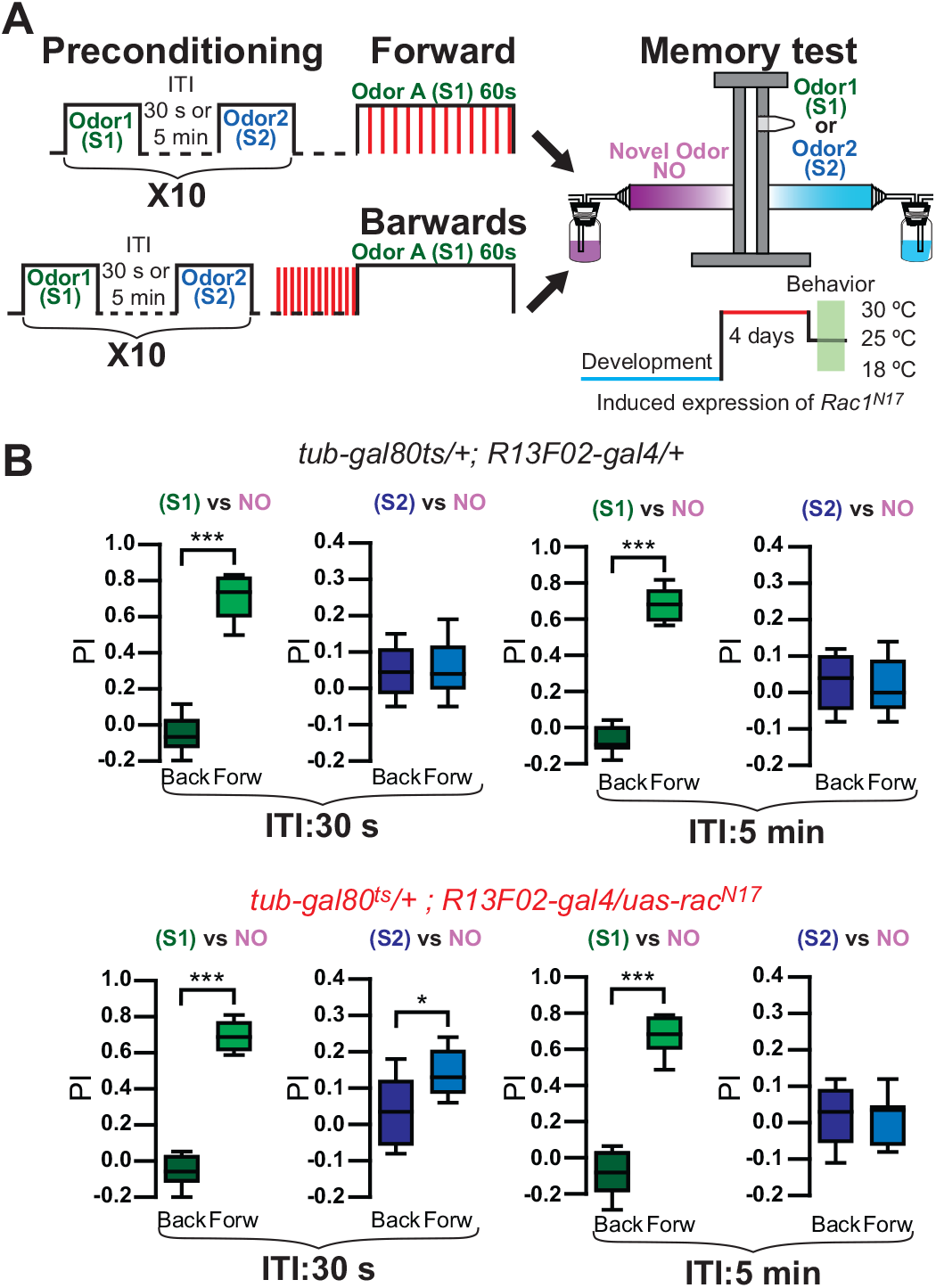
*Rac1* inhibition extends ISI for S1/S2 association for behavioral expression of sensory preconditioning. (A) Diagram of experimental setup (odor schedule): Flies were preconditioned by ten repeated presentations of S1/S2 odor pairs with either 30 s or 5 min ISI. Later flies were aversively trained to MCH (S1) by pairing one minute odor presentation along 12 90V, 1.25 s shocks. Right after training memory was tested in a T-maze by presenting either S1 vs a novel odor (NO) or S2 vs a NO. Performance index (PI) were compared to flies trained using backwards conditioning. B) Memory to S1 was significantly different from flies trained with backwards conditioning, for both control and flies expressing *Rac1*^*N17*^ in KC using a 30 s ISI. Non-parametric Mann-Whitney test *p<0*.*0002, n=8*. Similarly, memory to S2 in flies expressing *Rac1*^*N17*^ was significantly different from flies trained with backwards conditioning. Non-parametric Mann-Whitney test *p=0*.*0191, n=8*. In contrast, Memory to S2 was not significantly different in control flies. Non-parametric Mann-Whitney test *p>0*.*999, n=8*. Memory to S1 was significantly different from flies trained with backwards conditioning, for both control and flies expressing *Rac1*^*N17*^ in KC using a 5 min s ISI. Non-parametric Mann-Whitney test *p<0*.*0002, n=8*. In contrast, memory to S2 in both control and flies expressing *Rac1*^*N17*^ was not significantly different from flies trained with backwards conditioning. Non-parametric Mann-Whitney test *p≥0*.*7756, n=8*.

Taken all together, these results indicate that flies can infer value to non-paired odor based on the previous associative structure between odors and inhibition of *Rac1* by expression of its dominant-negative form in KC lengthens the time of olfactory “sensory buffer” allowing linking of neutral odors presented in sequence even when separated by at least 30 s.

## Material and Methods

### *Drosophila* husbandry

Flies were cultured on standard medium at room temperature. Crosses, unless otherwise stated, were kept at 25°C and 70% relative humidity with a 12 hr light-dark cycle. The drivers used in this study include *MB112C-splitgal4* ^18^, *R12G04-lexA*^59^, and *R13F02-gal4* ^59^. Drivers expression was verified by immunohistochemistry. Additional transgene stocks include d *uas-GCaMP6f* ^60^, *lexAop-GCaMP6f* ^60^, *uas-rac*^*N17* 61^, and *tub-gal80*^*ts* 20^. When Target system was used to restrict expression of transgene to adult animals, flies crosses were kept at 18 ºC during development. After eclosion 1-2 day old flies were transfer to 30 ºC for 4 days for the induction of expression. Flies were then transfer to 25 ºC 1 h before imaging or behavior. Control flies were subjected to exactly same protocol but they did not contain the UAS transgene. Additional controls were performed by keeping the crosses at 18 ºC during all time; these flies were then transferred to 25 ºC 1 h before imaging.

### *In vivo* calcium imaging

For measuring calcium responses with conditioning, odor or shock delivery, we processed flies as previously described with some modifications ^55,62^. Briefly, a single fly was aspirated without anesthesia into a narrow slot the width of a fly in a custom-designed recording chamber. The head was immobilized by gluing the eyes to the chamber using melted myristic acid and the proboscis fixed to reduce movements. A small, square section of dorsal cuticle was removed from the head to allow optical access to the brain. Fresh saline (103 mM NaCl, 3 mM KCl, 5 mM HEPES, 1.5 mM CaCl_2_, MgCl_2_, 26 mM NaHCO_3_, 1 mM NaH_2_PO_4_, 10 mM trehalose, 7 mM sucrose, and 10 mM glucose [pH 7.2]) was perfused immediately across the brain to prevent desiccation and ensure the health of the fly. Using a 20X water-immersion objective and a Leica TCS SP8 II confocal microscope with a 488 nm argon laser, we imaged the MBON-γ1pedc>α/β neuron for 2 min, during which stimuli was delivered starting at 30 s after imaging initiation. We used one HyD channel (510-550 nm) to detect *GCaMP6f* fluorescence.

### Odor and Shock Presentation

To deliver odors to flies under the microscope, a small stream of air (500 ml/min) was diverted (via solenoids) from flowing through a clean 20 ml glass vial to instead flow through a 20 ml glass vial containing a 0.5 µl drop of pure odorant. This air stream was then serial diluted into a larger air stream (1500 ml/min) before traveling through Teflon tubing (∼2.5 mm diameter) to reach the fly. To deliver shocks to flies under the microscope, a custom shock platform was made from shock grids used in standard olfactory memory assays that consist of alternating +/-charged copper strips attached to an epoxy sheet. To simulate shock exposure given during the standard olfactory memory assay, the surface of the shock platform was positioned so that all 6 legs are touching but the fly could temporarily break contact by moving its legs. Both solenoids for odors control and Grass stimulator for shock presentation were controlled by Arduino microcontroller (Arduino Uno) with custom-made programs.

### Training under microscope programs

The regular training protocol followed in most experiments on the paper, flies were presented to preconditioning odors with 5 s odor 1 followed by variable ISI where clean air was presented and by 5 s of a second odor (non-associated odor) pre-training (MCH and OCT or PA and EL). Five min after these preconditioning recordings, flies were trained under the microscope by simultaneous presentation of a single 20 s odor pulse and four 90V, 1.25 s electric shocks (5s inter shock interval). Five minutes after training, post-conditioning odor responses were recorded similarly to pre-responses. For control purposes, flies were trained with backwards training in which electric pulses were presented right before the onset of odor delivery.

### Behavior

Two to five day old flies were used for all behavior experiments. Standard aversive olfactory conditioning experiments were performed as described ^63^ with some modifications. Because we needed a precise control of odor delivery, this was achieved automatically using an Arduino microcontroller with custom-made programs to control a pair of solenoids. Briefly, a group of ∼60 flies were loaded into a training tube where they received a preconditional stimuli as indicated in each experiment. After preconditioning, flies were trained by 1 min of an odor paired with 12 pulses of 90V electric shock (S1). We used 3-octanol (OCT), 4-methylcyclohexanol (MCH), Penthyl acetate (PA) and Ethyl lactate (EL) as standard odorants. To measure memory, we transferred the flies into a T-maze where they were allowed 2 min to choose between the two odors either S1 or S2 and a novel, never experienced odor.

### Quantification and statistical analysis

Fluorescence was acquired from a region of interest (ROI) drawn around the axon tract of MBON-γ1pedc>α/β. Baseline was calculated using a Matlab code as the mean fluorescence across the 5 s before each odor presentation. This baseline was then used to calculate %ΔF/Fo for the complete recording. Bargraphs represent distribution of %ΔF/Fo responses across the 5 s of odor presentation. Solid lines in fluorescence traces represent mean %ΔF/Fo ± SE (shaded area) across the odor responses.

Statistics were performed using Prism 5 (Graphpad). All tests were two tailed and confidence levels were set at α=0.05. The figure legends present the *p* values and comparisons made for each experiment. Unless otherwise stated, non-parametric tests were used for all imaging and behavioral data.

## AUTHOR CONTRIBUTIONS

IC-S design of the study. AP, JC-M, and IC-S performed experiments. IC-S analyzed the data, interpreted the results and wrote an initial draft of the manuscript. AP and IC-S contributed to the final version of the manuscript.

## Acknowledgements

This work was supported by grants R21 MH117485-01A1 from the NIMH to IC-S, GU pilot research grant award ID162838 to IC-S.

